# Trade-offs between phage resistance and conjugative ability shape the ecological and evolutionary response of a multidrug resistance plasmid to plasmid-dependent phage

**DOI:** 10.1101/2025.10.08.681082

**Authors:** Daniel Cazares, Eliza Rayner, Adrian Cazares, Wendy Figueroa, David Goulding, Samuel Greenrod, Adam J. Mulkern, Michelle Yin, Tao He, Nick Thomson, Michael A. Brockhurst, R. Craig MacLean

**Author notes:** equal author contributions (Michael A. Brockhurst, R. Craig MacLean).

## Abstract

Phage therapy is a promising alternative to antibiotics to treat multidrug resistant infections. Plasmid dependent phages (PDPs) are particularly attractive as therapeutics because they can both kill targeted pathogen cells, whilst also potentially preventing the further spread of antibiotic resistance genes encoded by plasmids. However, we lack experimental studies of the ecological and evolutionary response of multidrug resistance plasmids against plasmid dependent phage treatment under ecologically relevant scenarios allowing plasmid conjugation. We experimentally evolved populations of E. coli carrying the multidrug resistance RP4 plasmid with the PRD1 PDP under conditions where conjugation was associated with either strong or weak benefits. When opportunities for conjugation were rare, PRD1 only transiently suppressed the conjugative plasmid population due to the rapid evolution of PRD1 resistant plasmids that lacked conjugative ability. Increasing the ecological opportunity for conjugation enhanced plasmid suppression and delayed the evolution of PRD1 resistant plasmids. PRD1 resistance was associated with reduced conjugative ability, but this trade-off was complex due to the heterogeneous impacts of resistance mutations on pilus production and conjugative ability. Mutations and IS element insertions in conjugation genes caused a wide range of PRD1 resistance phenotypes, ranging from complete resistance (*virB4*) to partial resistance (*trbB*, *trbL*). Bioinformatic analysis of publicly available IncP plasmid sequences showed that truncated variants of VirB4 protein are common in natural populations, suggesting that plasmid-dependent phages are an important selective pressure in microbial communities. Our results demonstrate an evolutionary trade-off between conjugative ability and phage resistance that cannot be easily circumvented by plasmids. Targeting multidrug resistance plasmids with PDPs is likely to drive loss of conjugation limiting the transfer of antibiotic resistance genes in bacterial communities.

## Introduction

The acquisition of conjugative plasmids carrying dedicated antibiotic resistance genes has played a key role in the evolution of antibiotic resistance in pathogenic bacteria^1,2^. Plasmids carrying antimicrobial resistance (AMR) genes tend to be associated with small fitness costs^3^, and conjugation enhances the ability of plasmids to persist in the absence of antibiotic use^4–6^, especially in communities ^7–10^. Given the inherent stability of plasmids^11–14^ it is important to develop active interventions to combat plasmid-mediated AMR^6,15–18^. This approach is likely to be particularly relevant for AMR plasmids that are distributed across connected bacterial host species and ecological niches (i.e. human, environment, agriculture) as interventions that target only one niche or host species may have limited impact on the prevalence of the plasmid^19–21^.

Plasmid-dependent phages (PDPs) use the conjugative pilus of plasmids as a receptor to infect bacterial cells^22–26^. PDPs are able to infect a wide range of bacterial hosts^27^, making them a promising tool to combat AMR plasmids that are widely disseminated across bacterial strains and/or species^28,29^ in bacterial communities that represent important reservoirs of antibiotic resistance, such as the gut microbiome^22,30,31^. A key challenge of using phage to combat AMR is that resistance can rapidly evolve, potentially compromising the efficacy of phage treatment^32^. For example, plasmids can adapt to PDPs through mutations that lead to a loss of the ability for conjugation^31^. In the short term, the loss of conjugative genes is likely to lead to a decreased ability of AMR plasmids to spread and persist in microbial communities, and the loss of conjugative ability is associated with an increased extinction probability of plasmid lineages over the long term^33^.

Evolutionary responses to PDPs have been studied by exposing monocultures of plasmid-carrying bacteria to phage. However, the lack of potential plasmid recipients in these experiments creates a scenario where conjugative ability is associated with very weak benefits^34,35^. Metabolic costs of conjugation often drive the evolution of non-conjugative plasmids under these conditions, even in the absence of phage pressure^36–38^. Given that trade-offs are likely to exist between conjugative ability and PDP resistance, we reasoned that increasing the potential for conjugative plasmid transfer reduces the strength of selection for PDP resistance and enhances the suppressive effect of PDPs on plasmid populations. To test our hypothesis, we challenged populations of *E. coli* carrying the RP4 plasmid with the PRD1 phage under conditions where opportunities for conjugation were either weak or strong by manipulating the rate of influx of plasmid-free immigrant cells into bacterial populations, a manipulation first used in a seminal paper by Dimitriu and colleagues^34^. We quantified the population dynamics of bacteria, plasmids and phage to understand the ecological consequences of this manipulation and we then used extensive phenotyping and genome sequencing to study the evolutionary response to phage under variable immigration treatments.

### Results Immigration enhances the suppressive effect of phage

To study the impact of PDP treatment, we established co-cultures of *E. coli* containing a plasmid-carrying strain (MG1655:RP4) and a plasmid-free strain (J53) that acted as a potential RP4 plasmid recipient. To manipulate the opportunity for conjugative transfer of the plasmid, populations were propagated by daily serial passaging under variable levels of immigration of plasmid-free J53 cells (0% immigration or 50% immigration). Bacteria and phage dynamics were tracked every 24 hours for 12 days, which corresponds to approximately 40 bacterial generations (Figure 1).

**Figure 1:**
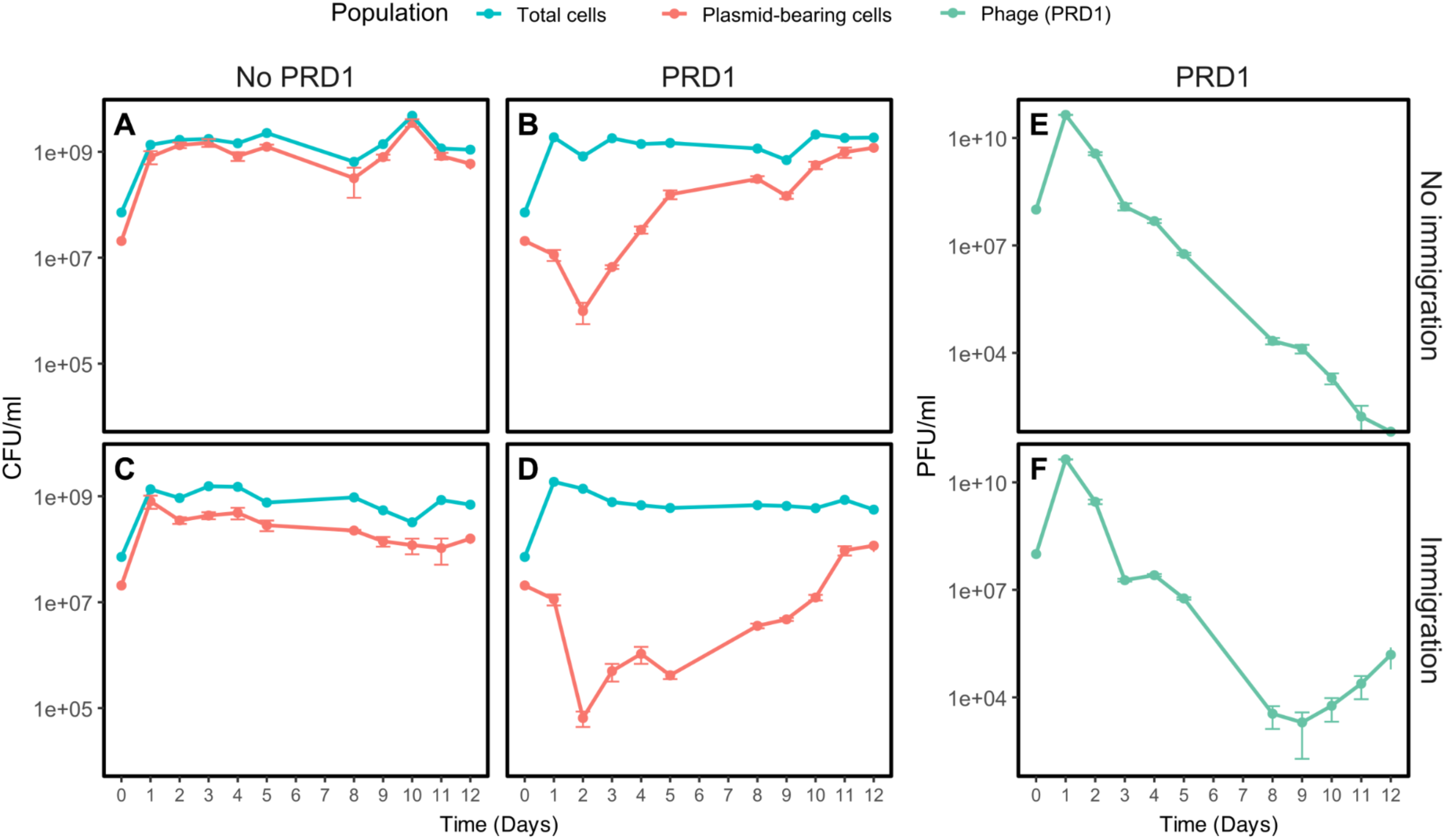
Immigration enhances the suppressive effect of phage. Plots show the mean total bacterial density, plasmid density, and phage density over time (+/- s.e; n=3). Panels correspond to the different conditions tested. **A)** No-phage, no-immigration, **B)** PRD1 treatment, no-immigration, **C)** No-phage with immigration, **D)** PRD1 treatment with immigration, while **E)** and **F)** correspond to the phage when immigration was absent or present, respectively. Treatment with phage suppressed the plasmid (main effect phage P=1.94E-05), and this effect was enhanced by continual immigration of plasmid free cells (phage*immigration interaction P=2.32E-03) as assessed with repeated measures ANOVA (supplementary Table 1). Immigration did not have an impact on phage density, as assessed by repeated measures ANOVA (P=.394 supplementary Table 2).

PRD1 treatment led to the rapid suppression of the RP4 plasmid, but this effect was transient, and plasmid densities eventually recovered to match phage-free controls. Immigration increased the short-term impact of phage treatment and slowed the long-term recovery of the plasmid, supporting our hypothesis that the efficacy of PDP treatment is greatest when opportunities for conjugation are plentiful. If immigration enhances the opportunity for conjugation, then it is reasonable to expect that immigration may lead to greater levels of phage replication and higher phage titers. However, phage densities were not significantly impacted by immigration, implying that the underlying strength of selection for phage resistance did not depend on immigration.

### PRD1 suppresses plasmids by killing donor cells and preventing plasmid transfer

To better understand the suppressive effects of PRD1 treatment, we assessed the dynamics of plasmid donors, recipients, and transconjugants (Figure 2, A-D, Supplementary data 1). Treatment with PRD1 was associated with lower densities of both donor cells and transconjugants. However, donors were typically 100x more abundant than transconjugants across all treatments, implying that the suppressive effect of PRD1 treatment on the RP4 plasmid (i.e. Figure 1) was driven by the killing of plasmid-carrying donor cells. As expected from Figure 1, phage led to transient suppression of donor cells in populations without immigration, whereas immigration was associated with more sustained suppression of donors by PRD1 (Figure 2E). In contrast, treatment with phage led to sustained suppression of transconjugants, independent of immigration rate (Figure 2F), highlighting the ability of PRD1 to prevent the transfer of plasmids between bacterial strains.

**Figure 2.**
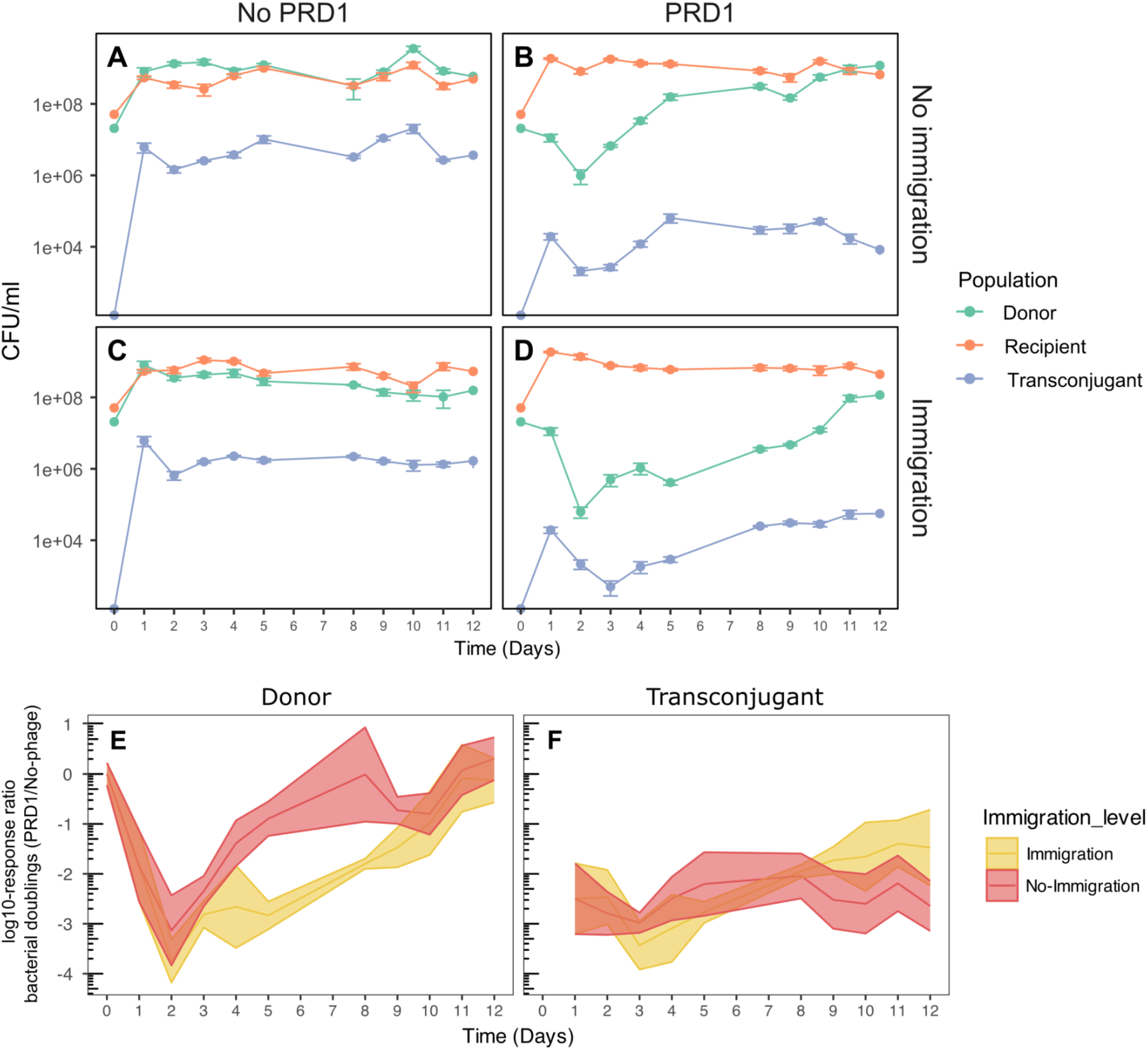
PRD1 treatment kills plasmid donor cells and prevents plasmid transfer across strains. Panels **A**-**D** show the mean density of plasmid donors, recipients and transconjugants over time (+/- s.e.; n=3). The ribbon plots in panels **E** and **F** show the log10-response ratio of daily doublings by the donor strain (E) and transconjugants (F) in PRD1-treated cultures relative to untreated controls. Lines show the estimated response ratio and the borders of the ribbon show the 95% confidence intervals in the response ratio.

### Immigration slows the evolution of phage resistance

The rapid decline in phage abundance (Figure 1) and recovery of the plasmid-bearing donor strain (Figures 1 and 2) suggest that resistance to PRD1 evolved. To test this idea, we isolated 360 clones from phage-evolving populations and 220 clones from phage-untreated samples and tested their susceptibility to the ancestral PRD1 phage. Three distinct phenotypes were detected, including phage susceptibility (clear PRD1 plaques), partial PRD1 resistance (cloudy plaques) and complete resistance (no plaques) (Figure 3A).

**Figure 3.**
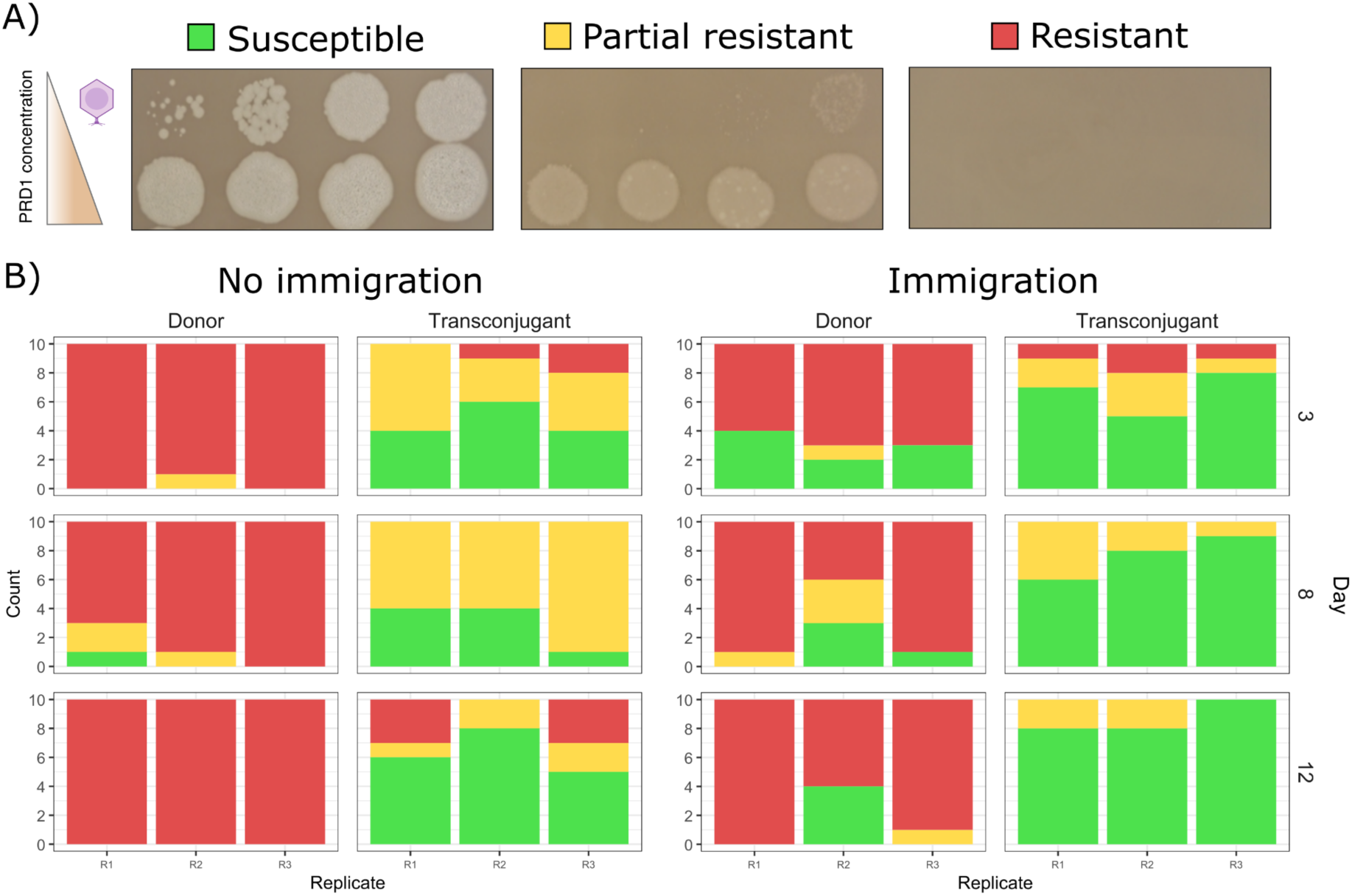
Distribution of phage resistance in plasmid populations. **A)** Representative results of PRD1 dilution spotting assays (10⁻⁸ to 10⁻¹) on lawns of isolates from the evolution experiments, classified as resistant (red), partial resistant (yellow), or susceptible (green). **B)** Distribution of these phenotypes across donor and transconjugant plasmid host populations for the 3 independent replicates of the evolution experiments under no-immigration and immigration conditions at days 3, 8, and 12. The proportion of phage resistant isolates (i.e. either full or partial resistance) in the donor population was higher in the absence of immigration at day 3 (Odds ratio=infinity, P=.0019), but not at day 8 (Odds ratio=4.46, P=.353) or day 12 (Odds ratio=infinity, P=.112).

The donor strain rapidly evolved resistance to PRD1 in the absence of immigration, with >95% of fully resistant isolates by day 3 of the experiment. Immigration delayed the evolution of phage resistance in the donor population, which is consistent with our hypothesis that opportunities for conjugation reduce the strength of selection for PDP resistance. Phage suppressed minority transconjugant sub-populations throughout the experiment, suggesting that sensitivity to phage remained high in transconjugants compared to donors. Consistent with this idea, the frequency of resistance remained low in transconjugants across the entire experiment (Figure 3B). Interestingly, phage resistance in transconjugants was usually partial, whereas the complete resistance phenotype was associated with donors.

In summary, isolate phenotyping revealed that resistance was high in donors compared to transconjugants, and more prevalent in treatments without immigration. To further test these associations, we carried out a side experiment where we re- introduced a high density of PRD1 to populations at day 7. Consistent with our phenotyping data, the suppressive effects of re-introducing PRD1 were most evident in the immigration treatment and on the transconjugants (Supplementary figure 2).

### Mutations in the conjugative pilus drive phage resistance evolution

To investigate evolutionary responses to PRD1 we sequenced endpoint populations and tested for the evolution of the RP4 plasmid by quantifying the abundance of novel mutations present at a frequency of >10% in the plasmid population. By this stage, plasmid populations were largely dominated by the donor strain, which displayed high levels of PRD1 resistance. PRD1 treatment was associated with parallel evolution across populations in the Tra2 region, which encodes the structural components of the type IV secretion system (T4SS), also known as the conjugative pilus. Most high-frequency mutations were SNPs or indels in *trbI*, *trbJ*, and *trbE*, reaching population frequencies of 30–77% (Figure 4, panels B and D). By contrast, in treatments without PRD1, no mutations in Tra2 exceeded 20% frequency, indicating that selection for altered conjugative ability is weak in the absence of phage (Figure 3, panels A and C). Low-frequency polymorphisms (10–20%) were also detected in other plasmid regions, including *traA* and *oriV*.

**Figure 4.**
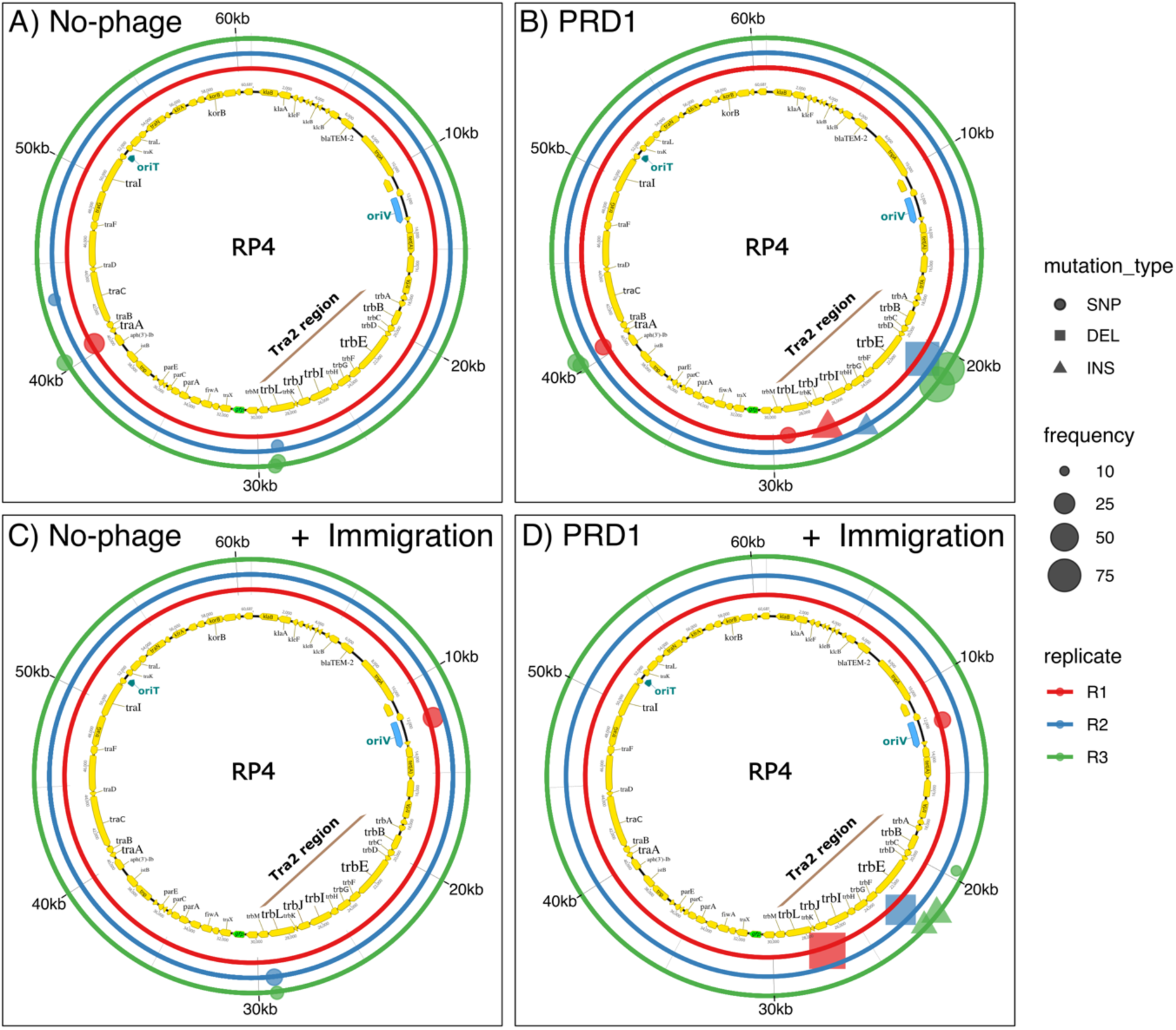
Plasmid population evolutionary responses to phage PRD1. Panels **A–D** show the genomic distribution and frequency of RP4 mutations detected at the endpoints of the evolution experiments. The inner ring depicts the RP4 genomic map (with the Tra2 region highlighted, where most mutations clustered). The three outer coloured rings represent biological replicates for each condition. Mutation types are indicated by symbols with shapes (circles = SNPs, squares = <u>DEL</u>ETIONS, triangles = <u>INS</u>ERTIONS), with symbol size proportional to mutation frequency.

### Evolutionary trajectories to phage resistance and associated trade-offs

To better understand the evolutionary responses of RP4 to PRD1, we further characterized 54 clones that were chosen to represent the full range of observed phage susceptibility phenotypes. To test for a trade-off between PRD1 resistance and conjugative ability, we quantitatively measured the phage susceptibility and conjugative ability of this panel of clones (Figure 5A, Supplementary data 1). As expected, we found a trade-off between phage resistance, measured as efficiency of plaquing, and conjugative ability, measured as relative transconjugant density in phage-free mating experiments. However, it is clear from this assay that phage resistance and conjugative ability are not binary traits, highlighting the ability of selection to fine-tune these traits (Figure 5A, Supplementary figure 3).

**Figure 5.**
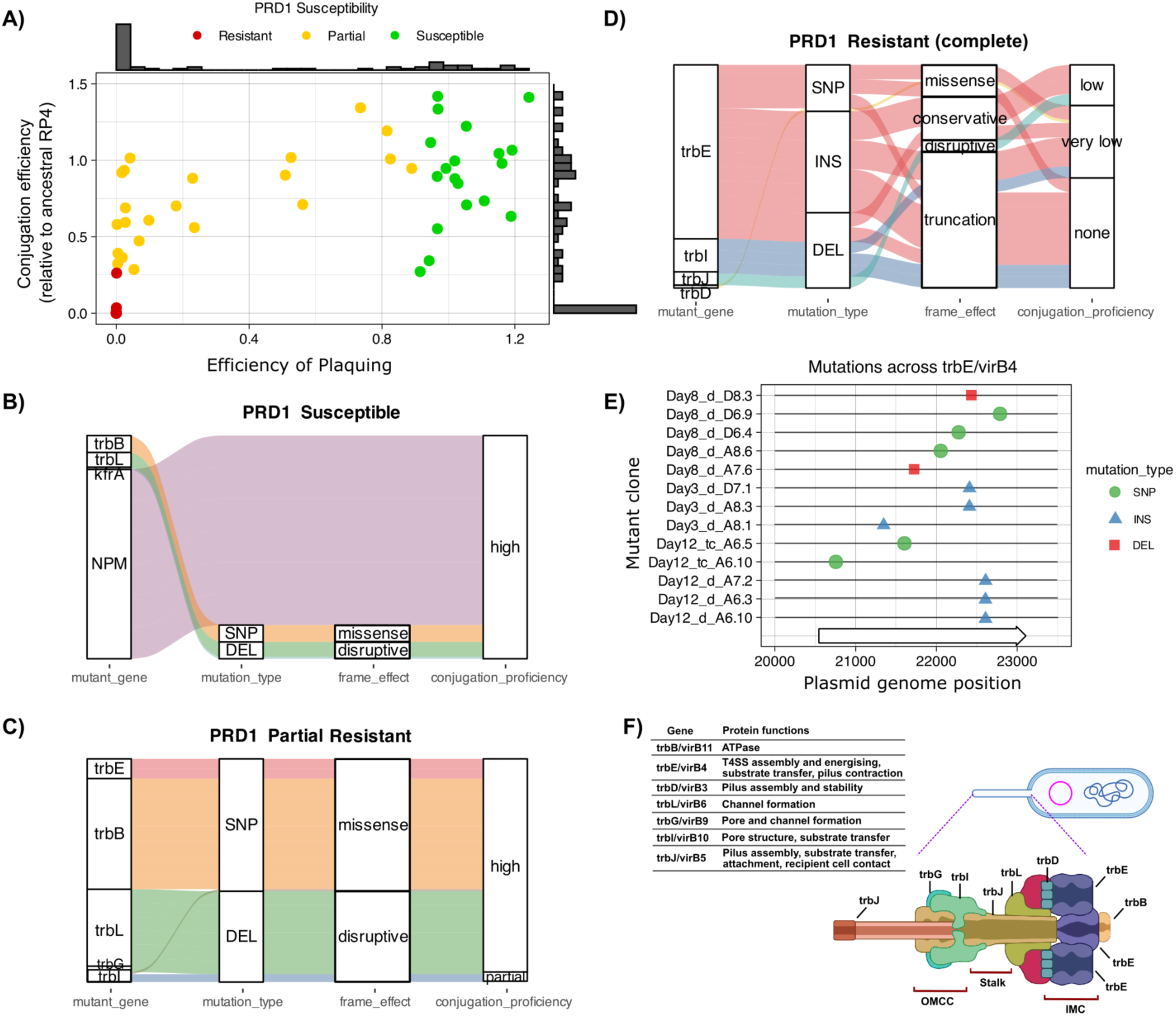
Phage resistance phenotypes and associated genotypes. **A)** Scatter plot of phage susceptibility (EOP, x-axis) versus conjugation efficiency (EOC, y-axis) for 65 RP4-carrying *E. coli* clones from the evolution experiments. Dots show mean values (3 replicates) and are color-coded by susceptibility: resistant (red), partial resistance (yellow), and susceptible (green). Values are relative to the ancestral RP4. Bar plots along each axis show the frequency distribution of phenotypes. **B-D** Panels show the links between plasmid mutations—including: the affected genes, mutation types (SNP, <u>del</u>etion, <u>ins</u>ertion), and their impact on the reading frame—and conjugation phenotypes. Phenotype and genotype associated with: **B)** PRD1-susceptible clones, **C)** Partial resistant clones, and **D)** Fully resistant clones. **E)** Distribution of mutations targeting *trbE/virB4*, the most frequently mutated gene among individual clones. **F)** Schematic of the type IV secretion system (T4SS) with regions showing the positions and predicted functions of proteins encoded by genes found to be mutated.

To understand the molecular basis of this trade-off, we sequenced plasmid carrying clones. As expected, phage-susceptible clones, which were recovered from no-phage and the phage treatments, were not associated with plasmid mutations (Supplementary data 2). Yet, some susceptible clones from the phage treatment were the exceptions, showing mutations in the genes *kfrA*, *trbB*, or *trbL*, but these mutations did not significantly affect conjugation (Figure 5B). Clones with a partial phage resistance phenotype were mainly associated with mutations in *trbB*, an ATPase that powers pilus retraction and extension, or *trbL*, which is involved in channel formation for the translocation of DNA. In all cases, these mutations were associated with a moderate decrease in conjugative ability, highlighting the capacity of plasmid RP4 to evolve increased resistance to PRD1 without completely sacrificing conjugative ability (Figure 5C, Supplementary figure 3).

Complete resistance to PRD1 was linked to mutations in *trbI*, *trbJ*, and *trbE*, matching the mutations identified by population sequencing (Figure 5D). We identified a wide diversity of mutations in *trbE*, including SNPs, INDELS, and insertions caused by the transposition of IS421 from the MG1655 chromosome to the plasmid, underscoring the role of IS elements in driving rapid evolution^39^. Despite differences in mutation type and position (Figure 5E), most were predicted to produce a truncated and misfolded TrbE protein (Supplementary figure 4).

TrbE plays key roles in the assembly and function of the T4SS, including inner membrane core complex (IMCC) assembly, energy supply, pilus contraction, and substrate translocation^44^ (Figure 5F). Consistent with this idea, mutations in *trbE* were associated with impaired conjugative ability. It is interesting to note, however, that *trbE* mutations had variable impacts on conjugative ability, and approximately 50% of clones with a mutation in *trbE* retained some level of conjugative ability, highlighting the complexity of the trade-off between conjugative ability and PRD1 resistance.

We also scanned these clones for chromosomal mutations to assess whether chromosomally encoded genes might contribute to the observed phenotypic outcomes. This analysis revealed that only 20% of the resistance clones had acquired chromosomal mutations. The affected genes were functionally diverse and scattered across different regions of the genome, and we could not identify any associations between these mutations and the phenotypes of interest (Supplementary table 3).

### Pilus is essential for phage-host infection

To investigate how the plasmid-encoded mutations associated with phage resistance affect the interaction with the bacterial host, we examined by transmission electron microscopy (TEM) *E. coli* cells carrying RP4 variants associated with either susceptibility or resistance to PRD1 and quantified phage-host binding. Representative resistant clones carried mutations in either *trbI* or *trbE*, while a susceptible clone lacking plasmid mutations served as a control. The plasmid-free strain J53 was included to evaluate the host’s background phage interaction level. In all cases, the plasmid-carrying isolates used in this assay lacked any chromosomal mutations.

TEM examination of 20 individual cells per clone (analysed in two rounds of independent experiments) revealed that PRD1 binds abundantly to the conjugative pilus—mostly to the sides of the pilus (Figure 6; clone without mutations at the top, Supplementary figure 5), a detail that remained unclear in earlier studies^45^. The susceptible clone displayed high phage binding, averaging 14 viral particles per cell, compared to resistant mutants, which bound far fewer phage—around 6 particles per cell for the *trbI* mutant and 2–3 for the *trbE* mutants (Figure 6). In line with the above, the *trbI* clone retained pilus remnants, whereas *trbE* mutants lacked these, likely explaining the higher binding in the former. Phage attachment to plasmid-free J53 cells was minimal (15× lower than the susceptible clone), consistent with weak, non-specific membrane interactions.

**Figure 6.**
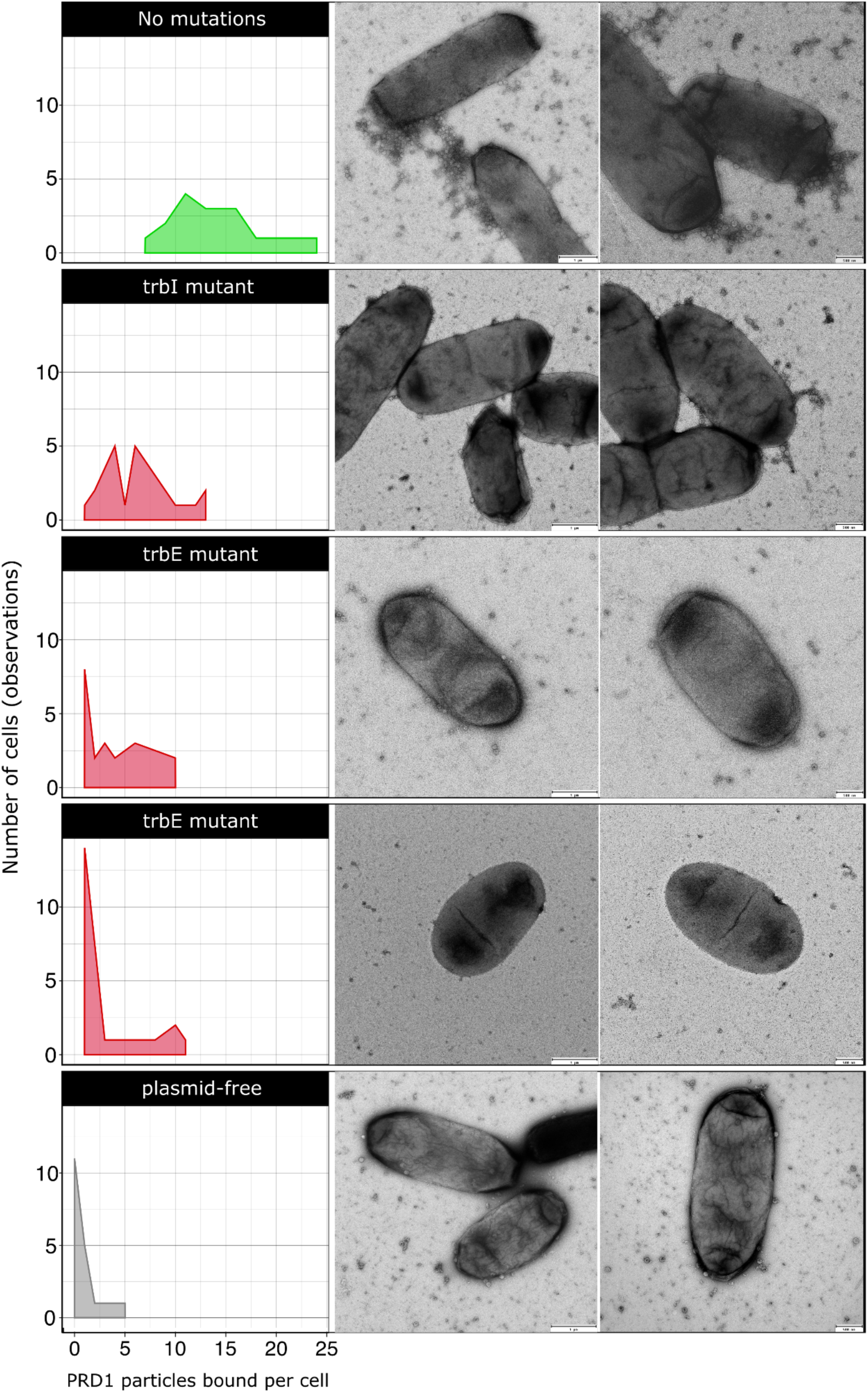
Bacterium–phage interactions via the conjugative pilus. TEM images show binding of phage PRD1 to *E. coli* cells carrying RP4 variants that are either phage-susceptible (top) or resistant (middle), with plasmid-free cells included as controls (bottom). Binding levels correlated with pilus presence: susceptible cells displayed abundant phage attachment, whereas resistant mutants with impaired pilus formation bound fewer viral particles. Frequency distribution plots at the left quantify PRD1 binding (x-axis) across cells (y-axis). The mutated gene of each variant is shown above the corresponding plot. Images on the left include a 100 nm scale bar, while those on the right include a 50 nm scale bar to provide higher magnification of the interaction phenotype.

We also observed that cells from the susceptible clone began to lyse as early as 10 minutes after exposure to the phage, which contrasted to the resistant mutants that remained intact throughout the examination. This rapid onset of cell bursting in the susceptible strain reinforces the phenotypic differences and highlights the protective effect conferred by the phage resistance-associated mutations.

### Truncations in the pilus assembly protein occur in natural IncP plasmid populations

Our experiments revealed that RP4 usually evolves resistance to PRD1 through mutations that truncate the protein TrbE, resulting in high levels of phage resistance and low conjugative ability. To determine whether similar genetic changes could be seen more broadly across publicly available IncP plasmid sequences, we investigated the diversity of TrbE/VirB4. A full-length alignment of 194 TrbE/VirB4 homologs— including the wild-type RP4 protein, representative mutant derivatives from our evolution experiment, and 182 homologs (≥60% sequence identity) retrieved from plasmid public archives—showed that protein length is generally well conserved, but also revealed notable variation, and a broad taxonomic distribution encompassing multiple classes (Figure 7A). Remarkably, we identified numerous truncated variants scattered across diverse branches of the TrbE/VirB4 phylogenetic tree—24 out of 182 (∼13%), excluding those from our evolution experiment, ranging from 67 to 846 aa. These truncated variants, lacking N-terminal or/and C-terminal sequences compared to RP4 TrbE/VirB4, were identified in diverse host genera, such as *Escherichia*, *Morganella* and *Pseudomonas*, and are present in a large proportion of plasmids (54%) encoding antimicrobial resistance (AMR) genes, including those conferring resistance to broad-spectrum beta-lactams or colistin (Supplementary figure 6, Supplementary data 2).

**Figure 7.**
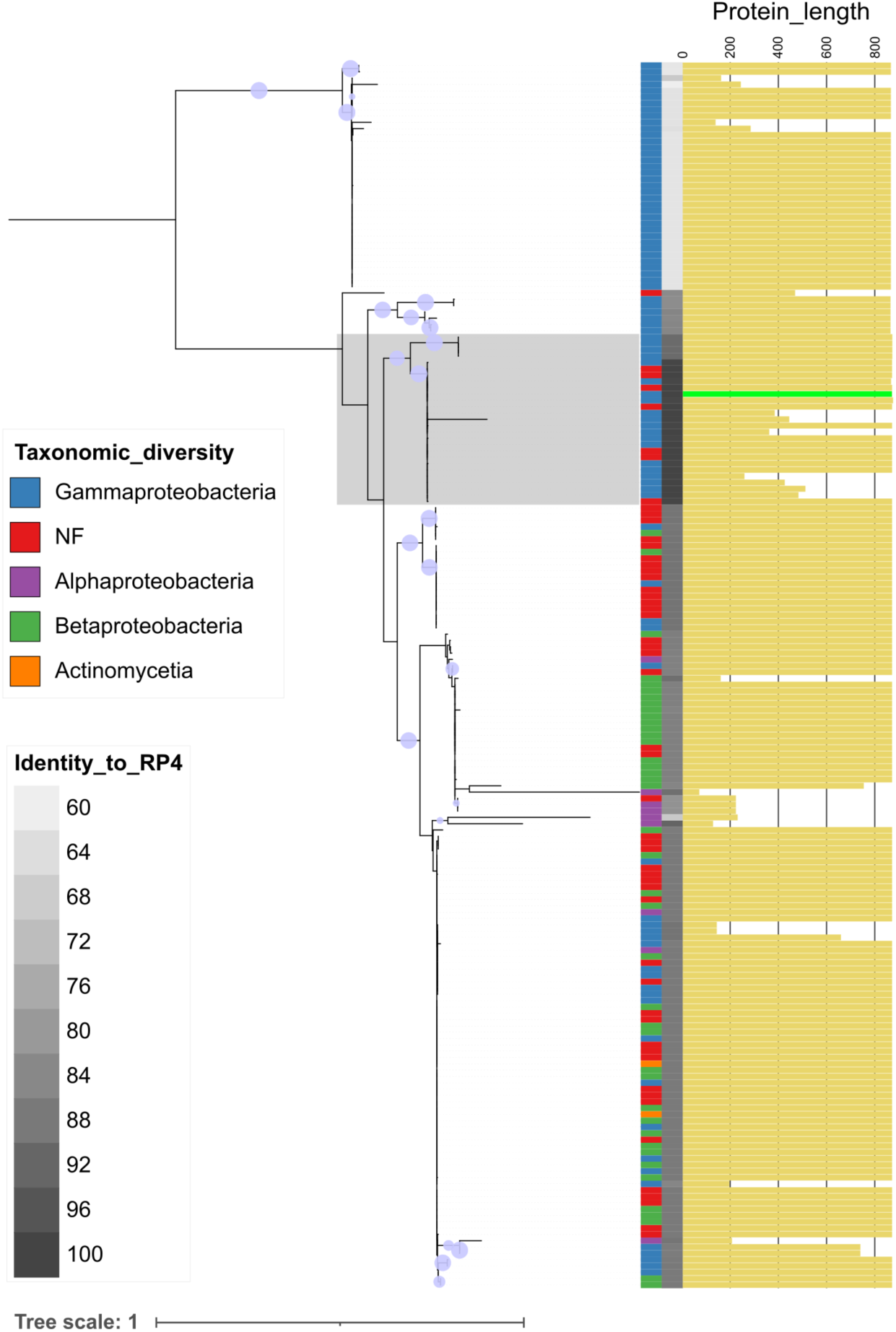
TrbE/VirB4 diversity in IncP plasmids. **A)** Phylogenetic tree of 194 TrbE/VirB4 homologs, including RP4 (length bar in green) and its evolved mutants— the region where they are distributed is highlighted. Protein length is shown by yellow bars (0–852 aa) on the right; shorter variants are present across multiple branches. Host taxonomic class is indicated by a color-coded bar, followed by sequence identity to RP4 (60–100%, grey gradient). Bootstrap support is marked by purple circles, where size is proportional to value (80–100). Tree scale indicates the number of substitutions per site under the model (LG+G4).

## Discussion

Plasmid-dependent phages (PDPs) are abundant and diverse^27^, but their impact on the evolution and ecology of plasmids remains poorly understood. In line with previous studies, we find that plasmids can evolve resistance to PRD1 through mutations that disable or impair the conjugative apparatus^31^. Although the loss of conjugative ability provides a simple solution to circumvent PDPs, mathematical models predict that conjugation plays a key role in plasmid maintenance^4,5,40^, and the loss of conjugation is associated with an elevated risk of plasmid lineage extinction^33^. Our work shows that PRD1 resistance evolves slowly under ecological conditions that create a benefit for conjugation (Figure 2, 3) because trade-offs between resistance and conjugative ability weaken selection for resistance (Figure 5). This, in turn, increases the suppressive effect of the PRD1 phage on the RP4 plasmid population (Figure 1, 2).

In addition to their suppressive effects on strains carrying plasmids, PDPs have the potential to block the horizontal transfer of plasmids between strains. As expected, PRD1 suppressed the transfer of the RP4 plasmid between strains of *E. coli*. Interestingly, levels of resistance to PRD1 in transconjugants remained low, despite the strong suppressive effect of phage on transconjugants. Because only conjugative plasmids can transfer, transconjugants must be biased towards conjugative plasmids compared to the donor population, and this inherent bias must help to explain the low levels of resistance in transconjugants. PRD1 pressure maintained transconjugants at a low population density (<10^4^ cells), leading to strong bottlenecks in the transconjugant population during passages to fresh medium (approximately 20-2000 cells transferred per passage). We speculate that low effective population size effectively prevented the evolution of RP4 resistance in the transconjugant sub- populations by limiting the potential for mutant plasmids to arise in the transconjugant sub-population and through the stochastic loss of rare PRD1 resistant transconjugant lineages.

Although a large number of genes are necessary for the formation of the RP4 conjugative pilus^41^, we only observed evolution in a sub-set of genes (*trbE, trbI, trbJ, trbD, trbB, trbL, trbG*) that have well described roles in pilus formation, DNA translocation and pilus retraction (Figure 5). Interestingly, some phage resistant mutants produced pili that provided a substrate for PRD1 attachment (Figure 6), suggesting that pilus retraction and substrate transfer are required for PRD1 to successfully infect cells. Mutations that truncate TrbE were the dominant mechanism of RP4 adaptation to PRD1 in our experiments. The assembly of the conjugative pilus is associated with substantial fitness costs^36–38^, and we speculate that TrbE truncation is a common evolutionary path to PRD1 resistance because the loss of TrbE prevents pilus formation, providing low cost PRD1 resistance. The *trbE* gene also presents a large mutational target compared to the other genes in the Tra region (2559 bp), which may also help to explain why this evolutionary trajectory is common. While we cannot entirely exclude the possibility that chromosomal genes may play a secondary role in modulating phage susceptibility or conjugative ability, the data indicate that their contribution was minor under the conditions of our experiment. Co-culturing bacteria and phage typically results in rapid co-evolution mediated by changes in bacterial cell surface components and phage tail fibre proteins^42,43^. PRD1 declined to very low phage densities (10^3^-10^4^ virions), making it very difficult to test for phage evolution in endpoint samples using either genomic or phenotypic assays. However, the continual decline in phage density over time suggests that PRD1 was not able to co-evolve to infect phage resistant bacteria, suggesting that the PRD1 may not be able to evolve the ability to infect bacteria that do not express functional conjugative pili.

Truncated variants of TrbE are common in IncP plasmids collected from diverse bacterial hosts and sources, including plasmids associated with antibiotic resistance genes (Figure 7). Although we cannot exclude the possibility that TrbE truncation reflects the outcome of selection associated with metabolic costs of pilus production, these results suggest that selection for resistance to PDPs may be widespread in nature. TrbE truncations in our study are associated with a dramatic loss of conjugative ability, suggesting that PRD1 may have played an important role naturally in preventing the dissemination of antibiotic resistance genes associated with IncP plasmids. One interesting avenue for research would be testing the hypothesis that the loss of conjugative ability in IncP plasmids is an evolutionary dead-end, as is suggested by large analyses of plasmids^33^.

The anthropogenic use of antibiotics over the past century has driven the evolution of conjugative plasmids carrying AMR genes^46^. The acquisition of these plasmids in pathogens is associated with stemming from the metabolic burden of plasmid gene expression and disruptions to core cellular processes^1,3^. Both theoretical and experimental studies show that conjugative transfer enables plasmids to persist in bacterial populations despite these costs, even under weak antibiotic pressure^5,6,8,9^. Our results indicate that PDPs impose a strong selective pressure that limits the conjugative ability of plasmids and constrains horizontal transfer of resistance genes within microbial communities. The growing discovery of PDPs highlights their potential as a biological toolkit to suppress diverse conjugative plasmids^47^. From this perspective, our findings suggest that targeting microbial communities that serve as hotspots for plasmid transfer using PDPs could simultaneously eliminate antibiotic-resistant pathogens and push plasmids to evolve into less mobile forms, limiting the onward dissemination of resistance.

## Methods

### Plasmid, bacterial, and phage strains

The multidrug resistance and promiscuous conjugative plasmid RP4 (which is a representative plasmid responsible for carbenicillin resistance outbreak in the UK during the 70s) was used for the evolution experiments. RP4 was tagged with the fluorescent marker GFP to facilitate plasmid detection. Briefly, the *gfp* gene, including promoter and terminator, was amplified from a synthetic template and ligated into linearised RP4 (digested with SnaBI and SbfI) using a Seamless Cloning reaction. The resulting RP4:GFP plasmid was introduced into *E. coli* DH5α by electroporation, then transferred by conjugation to *E. coli* MG1655, which served as the donor strain in evolution experiments (see Supplementary table 4 for details of the experimental conditions used). The plasmid-free strain *E. coli* J53 was used as the recipient. Selection was performed on LB agar with ampicillin (100 μg/mL) for MG1655(RP4:GFP), sodium azide (100 μg/mL) for J53, and both antibiotics for transconjugants. The plasmid-dependent phage PRD1 was used to investigate phage–plasmid interactions, and phage titers were determined by standard plaque assays.

### Experimental evolution setup

To study the evolutionary impact of PRD1 on RP4, we established serial passage systems of donor, recipient, and transconjugant *E. coli* populations, with or without phage. In a variant treatment, plasmid-free cells were added daily to promote conjugation (“immigration” condition). Cultures were inoculated with ∼1 × 10⁷ CFU/mL donors and ∼2 × 10⁷ CFU/mL recipients (1:2 ratio), and phage at ∼1 × 10⁸ PFU/mL (MOI ∼10). Independent cultures were maintained in triplicate in LB Lennox broth at 37 °C under static conditions. For the passages, cultures were diluted 1:10 into fresh medium each day. For immigration treatments, ∼2 × 10⁷ fresh recipients were added daily, while control treatments omitted phage (see Supplementary Figure 1 for experimental design).

### Tracking plasmid and phage dynamics

Population dynamics were monitored daily for 12 days by quantifying CFUs (donors, transconjugants, and recipients) and PFUs (PRD1). Briefly, culture aliquots were serially diluted (10⁻¹ to 10⁻⁸) in PBS with 5 mM sodium citrate (for bacteria) or in phage buffer (for PRD1). For CFU counts, 20 μL drops of each dilution were plated on selective plates to distinguish subpopulations; sodium citrate was included to minimise phage adsorption. For PFUs, dilutions were spotted on bacterial lawns of J53 carrying RP4. All spotting assays were performed in duplicate. Plates were incubated at 37 °C for 24 h. GFP fluorescence under blue light confirmed plasmid carriage in donor and transconjugant colonies.

To further test the extension of phage resistance during the evolution, PRD1 was reintroduced at high dose (∼1 × 10⁸) on day 7 in independent parallel populations of the experiments. This allowed us to assess whether PRD1 collapse was caused by the emergence of resistance rather than low phage concentrations, and whether immigration influenced resistance levels. Changes in plasmid host and phage densities were monitored as described above. Finally, the effect of immigration on plasmid–phage dynamics was quantified by calculating the response ratio of bacterial doublings (donors and transconjugants) in the presence versus absence of phage for both immigration and non-immigration conditions throughout the experiments.

### Phenotypic characterisation of evolved plasmid hosts

To assess the outcomes of the RP4–PRD1 interaction, we evaluated the frequency and degree of phage resistance in a library of 580 purified donor and transconjugant clones (360 from phage-treated populations and 200 from conditions without phage), sampled on days 3, 8, and 12 (30 colonies per population per time point). Phage susceptibility was determined by plaque assays and classified into three categories: (i) Susceptible – similar to the ancestral RP4 host; (ii) Partial resistance – producing cloudy plaques and showing resistance levels 5–1000-fold higher than the ancestral host; and (iii) Completely resistant – showing neither plaques nor lysis zones even under high phage densities (see Figure 3A).

A representative panel of 65 clones, covering all three phenotypes, was further tested for conjugation ability to evaluate the fitness cost of phage resistance. Conjugation assays were performed on LB Lennox plates using filter matings for 24 h at a donor-to-recipient ratio of 1:2. Cell densities were adjusted by OD600 measurements to add a similar number of cells for all assays. Depending on the donor strain, either *E. coli* J53 (when the donor was MG1655) or MG1655 chromosomally tagged with RFP (when the donor was J53) was used as the recipient to facilitate transconjugant selection. Both plaque and conjugation assays were performed in triplicate. Phage susceptibility was quantified as the efficiency of plaquing (EOP), calculated as the mean plaque number for the evolved clone divided by that of the ancestral host. Conjugation ability was quantified similarly as the efficiency of conjugation (EOC), using transconjugant counts instead of plaques.

### Genotypic characterisation of evolutionary outcomes

To investigate the evolutionary trajectories toward phage resistance, we sequenced and analysed both mixed endpoint populations from the evolution experiments and individual mutant clones displaying different phage susceptibility and conjugation phenotypes. Genomic DNA was extracted using the DNeasy Blood and Tissue Kit (QIAGEN), and plasmid DNA using the PureLink Quick Plasmid Miniprep Kit (invitrogen), both following the manufacturer’s protocols. Sequencing was performed on the Illumina platform (GENEWIZ UK Ltd). Illumina reads were quality-filtered (Trimmomatic) and used to generate plasmid de novo assemblies for individual clones with SPAdes^48^ v3.15.2.

Mutations were identified in both mixed populations and plasmid assemblies using Breseq^49^ v0.37.1 (https://github.com/barricklab/breseq). The option to predict polymorphic mutations was used to analyse the mixed population and default parameters were used for the individual clones, which were additionally scanned with Snippy v4.6.0 (https://github.com/tseemann/snippy). In both cases, the ancestral RP4 sequence was used as the reference to determine mutation positions and their predicted effects on proteins. Insertion of IS421 was identified through manual inspection of annotated sequences generated with BAKTA^50^ v1.11. Chromosomal mutations were assessed similarly, using the complete genomes of MG1655 (donors) or J53 (transconjugants) as references. Reference genomes were assembled using ONT long reads (PLASMIDSAURUS Ltd) and polished with Illumina reads (pypolca v0.2.0). The protein structures of *virB4/trbE* mutant variants were predicted with ColabFold^51^ v1.5.5 (https://github.com/sokrypton/ColabFold), combining AlphaFold2 and MMseqs2. Visualisation and structure sequence alignment comparison analysis was performed using ChimeraX^52^ (v1.9).

### Analysis of bacterium–phage interaction with electron microscopy

Transmission electron microscopy (TEM) was used to examine interactions between phage PRD1 and *E. coli* carrying RP4 derivative clones associated with either phage susceptibility or resistance. Clones with single mutations in *trbI* or *trbE* were selected as representative resistance genotypes, while a susceptible clone without plasmid mutations and plasmid-free *E. coli* J53 cells served as positive and negative controls, respectively.

Briefly, a single fresh colony grown overnight on LB Lennox agar plates was lifted onto a sterile loop and gently resuspended in an Eppendorf tube containing 100 µl double distilled water, turning it very slightly turbid. 5 µl of the cell suspension was added to the surface of a freshly glow-discharged carbon-coated Formvar EM grid and left for 30 seconds to allow settling of bacteria. The grid was gently blotted and 5 µl of neat PRD1 stock was then added and incubated for a further 30 seconds at room temperature. The grid was blotted again, negatively stained with 5 µL of 2% aqueous uranyl acetate and immediately blotted once more. Grids were analysed on a 120kV Tecnai Spirit Biotwin TEM at x20,000 magnification. Images were recorded using a F4.16 Tietz CCD. For each RP4 derivative clone, interactions were assessed in two independent experiments. Phage binding was quantified by counting the number of viral particles attached per cell from 20 independent observations.

For Scanning Electron Microscopy (SEM) of phage-pilus interaction, ten single fresh bacterial colonies (grown overnight on an LB Lennox agar plate) were each covered with 5 μl of neat PRD1 phage stock (∼1 × 10^11^), incubated for 30 seconds at room temperature, and gently rinsed three times with 0.1 M PBS. The colonies were then fixed on the plate with 1% paraformaldehyde and 2% glutaraldehyde in 0.1 M PBS for 1 hour, rinsed three times in 0.1 M sodium cacodylate, and post- fixed by layering 1% cacodylate-buffered osmium tetroxide and 1% aqueous thiocarbohydrazide, rinsing thoroughly in between. Individual blocks of agar 5mm x 5mm, containing single colonies, were cut from the plate and dehydrated through a graded ethanol series (30%, 50%, 70%, 95%, and 3 × 100%, 20 minutes each) and then dried in a Leica EM CPD 300. The samples were then coated with 6 nm of evaporated gold in a Leica ACE 600 sputter coater. Individual colonies were analysed and imaged in a Hitachi SU8030 scanning EM.

### TrbE/VirB4 distribution and phylogenetic analysis

IncP-group plasmids were identified from a global collection of plasmid genomes that integrates multiple plasmid databases^46^. Metadata for these plasmids (e.g. mobility, host, AMR gene content) was also retrieved as part of the collection. We searched the database via BLASTp^53^ using the protein sequence of TrbE/VirB4 from plasmid RP4 as a query. We restricted the matches to plasmids annotated as “IncP” according to the metadata available for the database (n=315). From these, we only kept plasmids with TrbE/VirB4 matches displaying at least 60% sequence similarity (n=182) for downstream analysis.

A full-length TrbE/VirB4 protein alignment combining the 182 sequences identified from public archives and 10 representative variants from our dataset was generated with muscle v3.8.1551^54^ using the “-sv -maxiters 2” options. The alignment was processed with trimal v1.4.1^55^ with the “-gt 0.9” parameter and a maximum-likelihood phylogeny was generated from it with raxml_tree from raxml-ng v1.1.0^56^ under the LG+G4 model using the following parameters: “--all --model LG+G4 --seed 12345 -- tree rand{10},pars{10} --bs-trees 100”. The tree was visualised in iTOL^57^ incorporating protein sequence length to illustrate the detected truncated variants.

## Data availability

Illumina data were deposited in GenBank. All sequence data from this study are associated with BioProject PRJNA1333424. SRA submission accessions SUB15661422 and SUB15661897 correspond to mixed population and individual clones data, respectively. BioSamples accessions can be found in the supplementary data set 2.

## Acknowledgements

This project received funding from UKRI (MR/W031361/1) and ICARS, under the umbrella of the JPIAMR-Joint Programming Initiative on Antimicrobial Resistance. This work was supported by UKRI Frontiers Grant (EP/Y031067/1) to RCM. This work was supported by funding from the Biotechnology and Biological Sciences Research Council (UKRI-BBSRC) [grant number BB/T008784/1] to ER. The authors would like to thank Matti Jalasvuori for kindly sharing PRD1. We thank Derek Pickard for technical advice on electron microscopy analysis.

## Author’s contributions

Conceptualisation: D.C., R.C.M., M.B., A.C.; Methodology: D.C., R.C.M., E.R., M.B., S.G., M.Y., A.M., T.H., W.F., A.C., D.G.; Data acquisition and analysis: D.C., E.R., R.C.M., S.G., A.M., W.F., A.C., D.G.; Funding acquisition: R.C.M., M.B., N.T.; Supervision: R.C.M., M.B.; Writing – original draft: D.C. and R.C.M., with input from other authors.; Writing – review & editing: All authors.

## References

1 MacLean, R. C. & San Millan, A. The evolution of antibiotic resistance. Science 365, 1082–1083 (2019). 10.1126/science.aax3879

2 Partridge, S. R., Kwong, S. M., Firth, N. & Jensen, S. O. Mobile Genetic Elements Associated with Antimicrobial Resistance. Clinical Microbiology Reviews 31 (2018). 10.1128/cmr.00088-17

3 Vogwill, T. & MacLean, R. C. The genetic basis of the fitness costs of antimicrobial resistance: a meta-analysis approach. Evolutionary Applications 8, 284–295 (2015). 10.1111/eva.12202

4 Lundquist, P. & Levin, B. TRANSITORY DEREPRESSION AND THE MAINTENANCE OF CONJUGATIVE PLASMIDS. GENETICS 113, 483–497 (1986).

5 Stewart, F. & Levin, B. POPULATION BIOLOGY OF BACTERIAL PLASMIDS - APRIORI CONDITIONS FOR EXISTENCE OF CONJUGATIONALLY TRANSMITTED FACTORS. GENETICS 87, 209–228 (1977).

6 Lopatkin, A. J. et al. Persistence and reversal of plasmid-mediated antibiotic resistance. Nature Communications 8 (2017). 10.1038/s41467-017-01532-1

7 Hall, J., Williams, D., Paterson, S., Harrison, E. & Brockhurst, M. Positive selection inhibits gene mobilization and transfer in soil bacterial communities. NATURE ECOLOGY & EVOLUTION 1, 1348–1353 (2017). 10.1038/s41559-017-0250-3

8 Hall, J., Wood, A., Harrison, E. & Brockhurst, M. Source-sink plasmid transfer dynamics maintain gene mobility in soil bacterial communities. PROCE1EDINGS OF THE NATIONAL ACADEMY OF SCIENCES OF THE UNITED STATES OF AMERICA 113, 8260–8265 (2016). 10.1073/pnas.1600974113

9 Alonso-del Valle, A., et al. Variability of plasmid fitness effects contributes to plasmid persistence in bacterial communities. Nature Communications 12 (2021). 10.1038/s41467-021-22849-y

10 León-Sampedro, R. et al. Pervasive transmission of a carbapenem resistance plasmid in the gut microbiota of hospitalized patients. NATURE MICROBIOLOGY 6, 606–+ (2021). 10.1038/s41564-021-00879-y

11 Yassour, M. et al. Natural history of the infant gut microbiome and impact of antibiotic treatment on bacterial strain diversity and stability. Sci. Transl. Med. 8 (2016). 10.1126/scitranslmed.aad0917

12 Ogunlana, L. et al. Regulatory fine-tuning of mcr-1 increases bacterial fitness and stabilises antibiotic resistance in agricultural settings. ISME JOURNAL 17, 2058– 2069 (2023). 10.1038/s41396-023-01509-7

13 He, W., Russel, J., Klincke, F., Nesme, J. & Sorensen, S. Insights into the ecology of the infant gut plasmidome. NATURE COMMUNICATIONS 15 (2024). 10.1038/s41467-024-51398-3

14 Kang, J. et al. Long-term ecological and evolutionary dynamics in the gut microbiomes of carbapenemase-producing Enterobacteriaceae colonized subjects. NATURE MICROBIOLOGY 7, 1516–+ (2022). 10.1038/s41564-022-01221-w

15 Lopez-Igual, R., Bernal-Bayard, J., Rodriguez-Paton, A., Ghigo, J. M. & Mazel, D. Engineered toxin-intein antimicrobials can selectively target and kill antibiotic-resistant bacteria in mixed populations. Nat Biotechnol 37, 755–760 (2019). 10.1038/s41587-019-0105-3

16 Lloyd, G., Stephens, E., Di Maio, A. & Thomas, C. Activation, incompatibility, and displacement of FIB replicons in E. coli. NUCLEIC ACIDS RESEARCH 53 (2025). 10.1093/nar/gkaf275

17 Lujan, S., Guogas, L., Ragonese, H., Matson, S. & Redinbo, M. Disrupting antibiotic resistance propagation by inhibiting the conjugative DNA relaxase. PROCEEDINGS OF THE NATIONAL ACADEMY OF SCIENCES OF THE UNITED STATES OF AMERICA 104, 12282–12287 (2007). 10.1073/pnas.0702760104

18 Getino, M. et al. Synthetic Fatty Acids Prevent Plasmid-Mediated Horizontal Gene Transfer. MBIO 6 (2015). 10.1128/mBio.01032-15

19 Shaw, L. P. et al. Niche and local geography shape the pangenome of wastewater- and livestock-associated Enterobacteriaceae. Sci. Adv. 7 (2021). 10.1126/sciadv.abe3868

20 Wang, Y. et al. Changes in colistin resistance and mcr-1 abundance in Escherichia coli of animal and human origins following the ban of colistin-positive additives in China: an epidemiological comparative study. Lancet Infect. Dis. 20, 1161–1171 (2020). 10.1016/s1473-3099(20)30149-3

21 Kumarasamy, K. K. et al. Emergence of a new antibiotic resistance mechanism in India, Pakistan, and the UK: a molecular, biological, and epidemiological study. Lancet Infect. Dis. 10, 597–602 (2010). 10.1016/s1473-3099(10)70143-2

22 Penttinen, R., Given, C. & Jalasvuori, M. Indirect Selection against Antibiotic Resistance via Specialized Plasmid-Dependent Bacteriophages. MICROORGANISMS 9 (2021). 10.3390/microorganisms9020280

23 Coetzee, J. N., Bradley, D. E., Dutoit, L. & Hedges, R. W. BACTERIOPHAGE-X-2 - A FILAMENTOUS PHAGE LYSING INCX-PLASMID-HARBOURING BACTERIAL STRAINS. Journal of General Microbiology 134, 2535–2541 (1988).

24 Bradley, D. E., Coetzee, J. N., Bothma, T. & Hedges, R. W. PHAGE-X - A PLASMID-DEPENDENT, BROAD HOST RANGE, FILAMENTOUS BACTERIAL-VIRUS. Journal of General Microbiology 126, 389–396 (1981).

25 Olsen, R., Siak, J. & Gray, R. CHARACTERISTICS OF PRD1, A PLASMID-DEPENDENT BROAD HOST RANGE DNA BACTERIOPHAGE. JOURNAL OF VIROLOGY 14, 689–699 (1974).

26 Parra, B. et al. Isolation and characterization of novel plasmid-dependent phages infecting bacteria carrying diverse conjugative plasmids. MICROBIOLOGY SPECTRUM 12 (2024). 10.1128/spectrum.02537-23

27 Quinones-Olvera, N. et al. Diverse and abundant phages exploit conjugative plasmids. NATURE COMMUNICATIONS 15 (2024). 10.1038/s41467-024-47416-z

28 Acman, M. et al. Role of mobile genetic elements in the global dissemination of the carbapenem resistance gene bla(NDM). Nature Communications 13 (2022). 10.1038/s41467-022-28819-2

29 Wang, R. B. et al. The global distribution and spread of the mobilized colistin resistance gene mcr-1. Nature Communications 9 (2018). 10.1038/s41467-018-03205-z

30 Colom, J. et al. Sex pilus specific bacteriophage to drive bacterial population towards antibiotic sensitivity. SCIENTIFIC REPORTS 9 (2019). 10.1038/s41598-019-48483-9

31 Jalasvuori, M., Friman, V., Nieminen, A., Bamford, J. & Buckling, A. Bacteriophage selection against a plasmid-encoded sex apparatus leads to the loss of antibiotic-resistance plasmids. BIOLOGY LETTERS 7, 902–905 (2011). 10.1098/rsbl.2011.0384

32 Pirnay, J. et al. Personalized bacteriophage therapy outcomes for 100 consecutive cases: a multicentre, multinational, retrospective observational study. NATURE MICROBIOLOGY 9, 1434–+ (2024). 10.1038/s41564-024-01705-x

33 Coluzzi, C., Garcillan-Barcia, M., de la Cruz, F. & Rocha, E. Evolution of Plasmid Mobility: Origin and Fate of Conjugative and Nonconjugative Plasmids. MOLECULAR BIOLOGY AND EVOLUTION 39 (2022). 10.1093/molbev/msac115

34 Dimitriu, T., Matthews, A. & Buckling, A. Increased copy number couples the evolution of plasmid horizontal transmission and plasmid-encoded antibiotic resistance. PROCEEDINGS OF THE NATIONAL ACADEMY OF SCIENCES OF THE UNITED STATES OF AMERICA 118 (2021). 10.1073/pnas.2107818118

35 Anderson, R. & May, R. COEVOLUTION OF HOSTS AND PARASITES. PARASITOLOGY 85, 411–426 (1982). 10.1017/S0031182000055360

36 Porse, A., Schonning, K., Munck, C. & Sommer, M. Survival and Evolution of a Large Multidrug Resistance Plasmid in New Clinical Bacterial Hosts. MOLECULAR BIOLOGY AND EVOLUTION 33, 2860–2873 (2016). 10.1093/molbev/msw163

37 Souque, C., Escudero, J. & MacLean, R. Off-Target Integron Activity Leads to Rapid Plasmid Compensatory Evolution in Response to Antibiotic Selection Pressure. MBIO 14 (2023). 10.1128/mbio.02537-22

38 Millan, A. S. & Maclean, R. C. Fitness Costs of Plasmids: a Limit to Plasmid Transmission. Microbiology Spectrum 5 (2017). 10.1128/microbiolspec.MTBP-0016-2017

39 Sastre-Dominguez, J. et al. Plasmid-encoded insertion sequences promote rapid adaptation in clinical enterobacteria. NATURE ECOLOGY & EVOLUTION 8 (2024). 10.1038/s41559-024-02523-4

40 Bergstrom, C., Lipsitch, M. & Levin, B. Natural selection, infectious transfer and the existence conditions for bacterial plasmids. GENETICS 155, 1505–1519 (2000).

41 Grahn, A., Haase, J., Lanka, E. & Bamford, D. Assembly of a functional phage PRD1 receptor depends on 11 genes of the IncP plasmid mating pair formation complex. JOURNAL OF BACTERIOLOGY 179, 4733–4740 (1997). 10.1128/jb.179.15.4733-4740.1997

42 Betts, A., Gray, C., Zelek, M., MacLean, R. C. & King, K. C. High parasite diversity accelerates host adaptation and diversification. Science 360, 907–911 (2018). 10.1126/science.aam9974

43 Paterson, S. et al. Antagonistic coevolution accelerates molecular evolution. Nature 464, 275–278 (2010). 10.1038/nature08798

44. Macé, K., Vadakkepat, A.K., Redzej, A. et al. Cryo-EM structure of a type IV secretion system. Nature 607, 191–196 (2022). 10.1038/s41586-022-04859-y

45. Kotilainen MM, Grahn AM, Bamford JK, Bamford DH.1993. Binding of an Escherichia coli double-stranded DNA virus PRD1 to a receptor coded by an IncP-type plasmid. J Bacteriol 175. 10.1128/jb.175.10.3089-3095.1993

46. Adrian Cazares et al., Pre- and postantibiotic epoch: The historical spread of antimicrobial resistance. Science 0,eadr1522 10.1126/science.adr1522

47. Cazares, D., Figueroa, W. & Cazares, A. Rediscovering plasmid-dependent phages. Nat Rev Microbiol 22, 670 (2024). 10.1038/s41579-024-01102-5

48. Prjibelski, A., Antipov, D., Meleshko, D., Lapidus, A., & Korobeynikov, A. (2020). Using SPAdes de novo assembler. Current Protocols in Bioinformatics, 70, e102. 10.1002/cpbi.102

49. Barrick, J., Yu, D., Yoon, S. et al. Genome evolution and adaptation in a long-term experiment with Escherichia coli. Nature 461, 1243–1247 (2009). 10.1038/nature08480

50. Schwengers O., Jelonek L., Dieckmann M. A., Beyvers S., Blom J., Goesmann A. (2021). Bakta: rapid and standardized annotation of bacterial genomes via alignment-free sequence identification. Microbial Genomics, 7(11). 10.1099/mgen.0.000685

51. Mirdita, M., Schütze, K., Moriwaki, Y. et al. ColabFold: making protein folding accessible to all. Nat Methods 19, 679–682 (2022). 10.1038/s41592-022-01488-1

52. Meng EC, Goddard TD, Pettersen EF, Couch GS, Pearson ZJ, Morris JH, et al. UCSF ChimeraX: Tools for structure building and analysis. Protein Science. 2023; 32(11):e4792. 10.1002/pro.4792

53. Altschul, S. F., Gish, W., Miller, W., Myers, E. W., & Lipman, D. J. (1990). Basic local alignment search tool. Journal of Molecular Biology, 215(3), 403–410. 10.1016/S0022-2836(05)80360-2

54. Robert C. Edgar, MUSCLE: multiple sequence alignment with high accuracy and high throughput, Nucleic Acids Research, Volume 32, Issue 5, 1 March 2004, Pages 1792–1797,. 10.1093/nar/gkh340

55. Salvador Capella-Gutiérrez, José M. Silla-Martínez, Toni Gabaldón, trimAl: a tool for automated alignment trimming in large-scale phylogenetic analyses, Bioinformatics, Volume 25, Issue 15, August 2009, Pages 1972–1973,. 10.1093/bioinformatics/btp348

56. Alexey M Kozlov, Diego Darriba, Tomáš Flouri, Benoit Morel, Alexandros Stamatakis, RAxML-NG: a fast, scalable and user-friendly tool for maximum likelihood phylogenetic inference, Bioinformatics, Volume 35, Issue 21, November 2019, Pages 4453–4455,. 10.1093/bioinformatics/btz305

57. Ivica Letunic, Peer Bork, Interactive Tree of Life (iTOL) v6: recent updates to the phylogenetic tree display and annotation tool, Nucleic Acids Research, Volume 52, Issue W1, 5 July 2024, Pages W78–W82,. 10.1093/nar/gkae268

